# EvoKEN: evolutionary knowledge extraction in networks

**DOI:** 10.1101/098285

**Authors:** Benjamin Linard, Ngoc Hoan Nguyen, Odile Lecompte, Olivier Poch, Julie D. Thompson

## Abstract

We introduce a multi-factorial, multi-level approach to build and explore evolutionary scenarios of complex protein networks. EvoKEN combines a unique formalism for integrating multiple types of data associated with network molecular components and knowledge extraction techniques for detecting cohesive/anomalous evolutionary processes. We analyzed known human pathway maps and identified perturbations or specializations at the local topology level that reveal important evolutionary and functional aspects of these cellular systems.

---

The dynamic molecular machinery underlying cellular systems is often represented by complex, hierarchical networks of interactions between the cell’s constituents, such as proteins, DNA, RNA and small molecules. Ultimately, phenotypic traits and diseases can be described in terms of the complex intracellular and intercellular networks that link tissue and organ systems^1,2^. The structures of these networks, including metabolic, signaling or transcription regulatory networks, often share similar features even in distantly related species^3^. Understanding the evolution of these networks is therefore essential to reconstruct the history of life, but also to better understand how the network structures correlate with the functioning of organisms at different granularity levels^4,5^.

Application of evolutionary based methods in complex networks is challenging^6^ and requires integration of multiple factors, such as gene spatial/temporal expression, protein sequence conservation, cellular localization signals, 3D structure, or binding/interaction sites^7^. In this context, we have developed an original formalism, called the evolutionary barcode or EvoluCode^8^, to allow the integration of different parameters (e.g. genome context, protein organization, conservation patterns) in a common framework and to summarize the evolutionary history of a gene that leads to its current state in a given organism (Supplementary Fig. 1). EvoluCode thus facilitates the application of formal data mining and knowledge extraction techniques in evolutionary analyses. We previously used this approach to barcode all human protein-coding genes using 10 evolutionary data types from 17 vertebrate proteomes. Our systematic comparison of the human barcodes revealed protein function-evolution relationships that could not be observed by using only one or two biological parameters, for example using only sequence conservation^8^.

Here, we introduce a unique protocol, called EvoKEN (Evolutionary Knowledge Extraction in Networks), that combines the EvoluCode formalism with knowledge extraction techniques, in order to study the evolution of genes in the context of their complex biological networks. We show how EvoKEN can be used at the pathway level to identify local topological motifs that have evolved cohesively and to highlight ‘outlier’ genes whose evolutionary history deviates from the local neighbors, suggesting different underlying evolutionary processes. We then extend our work to investigate unusual evolutionary scenarios at the inter-pathway or ‘cellular’ level.

Our protocol can be applied to any biological system, where we define a system as a set of genes implicated in a common process or phenomenon (genetic information processing, signal transduction, metabolism, disease response, etc.) and mapped onto a molecular network. We demonstrate the utility of our approach by constructing and exploring evolutionary scenarios for the complete set of human pathway maps in the Kyoto Encyclopedia of Genes and Genomes (KEGG) knowledge base^9^. First, we mapped our EvoluCodes for the human proteome to the KEGG maps, thus producing pathway-level evolutionary maps (Fig. 1) for a total of 248 biological systems (available at lbgi.igbmc.fr/barcodes). We then applied a knowledge extraction algorithm (Online Methods) on each individual map in order to estimate its evolutionary cohesiveness and to identify genes with anomalous, ‘outlier’ barcodes that might reflect unusual evolutionary pressures within the system. Here, we used the Local Outlier Factor (LOF)^10^, a powerful anomaly detection algorithm which is related to density-based clustering and is suitable for analyzing large-scale, multidimensional datasets where the underlying data distribution is unknown. The LOF method identified a total of 1147 outlier genes in 248 KEGG maps (lbgi.igbmc.fr/barcodes and Supplementary Fig. 2). The most cohesive pathways, i.e. those with the least outliers (Supplementary Fig. 3 and Supplementary Table 1), were typically involved in universal biological processes such as translation or cell growth/death, in line with previous observations^11^.

**Figure 1.**
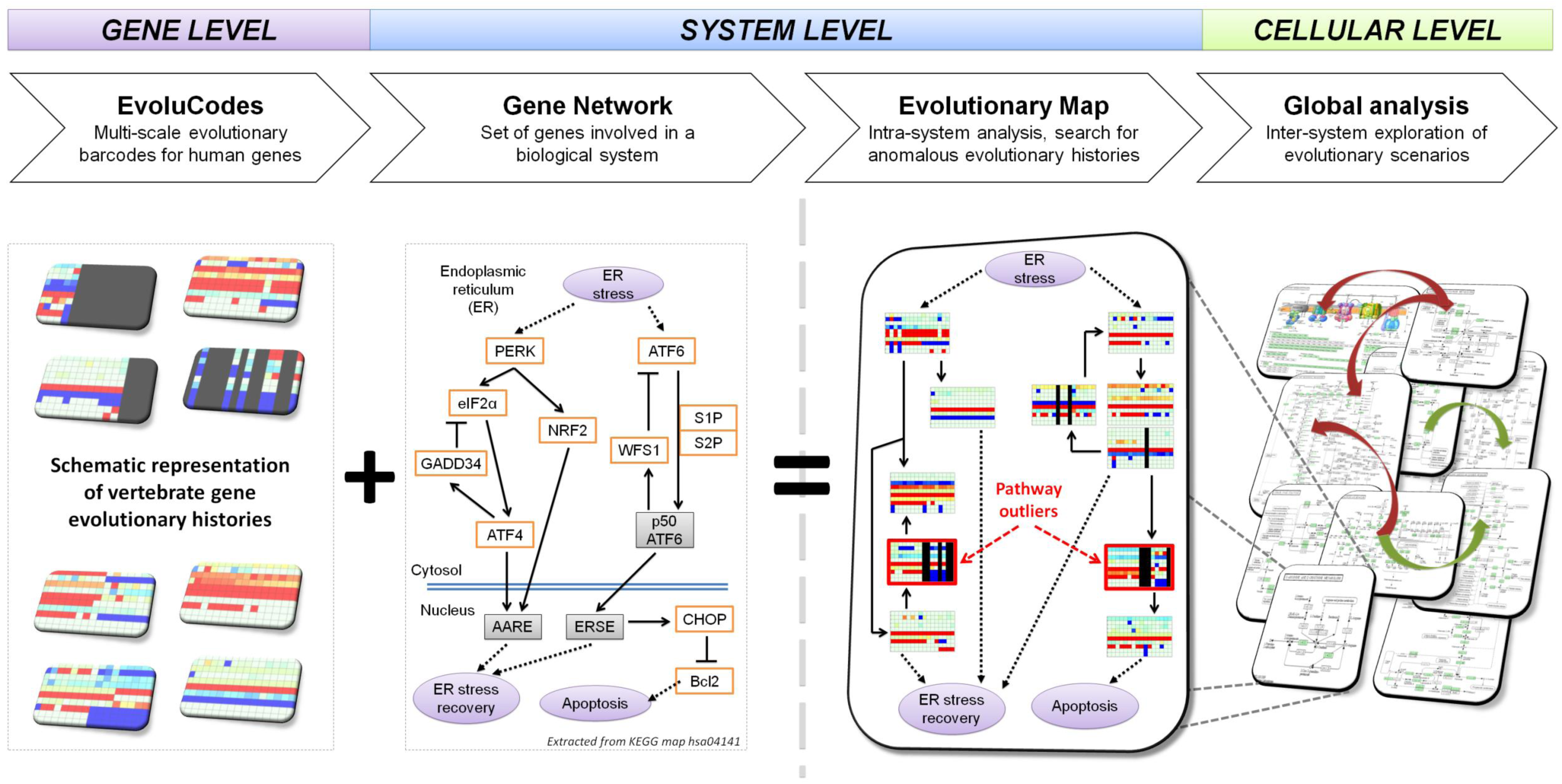
Framework to construct and explore multi-level evolutionary networks. Evolutionary barcodes, known as EvoluCodes, are assigned to individual genes and then mapped onto a known gene network, such as a KEGG pathway map. At the system level, the resulting evolutionary map provides a context for differentiating genes with 'cohesive' or 'outlier' (highlighted in red) evolutionary histories. At the cellular level, systems and inter-system crosstalk can be analyzed in terms of the cohesiveness of the underlying gene evolution.

To further investigate the biological significance of genes with anomalous evolutionary histories, we measured the correlations between the EvoKEN outliers and their local topology in the corresponding networks. We focused on the metabolic pathways in KEGG, where the nodes in the networks represent metabolites (substrates, products and intermediates) that are linked by a reaction, associated with one or more genes/proteins. Within these networks, we manually defined 6 classes of local topological motifs based on 2 key node properties, redundancy and connectivity (Fig. 2a and Online Methods). The outlier genes from 20 metabolic pathways were then assigned to the different topology classes (Fig. 2b). We found that the cohesiveness of a gene in its network context depends on the local topological structure: for instance, the smallest proportion of outliers was found at the nodes involved in linear paths in the networks, particularly in non-redundant paths (class F). In contrast, more outliers were found at the start/end points of a pathway (class D), and at the interface between pathways, so called ‘hubs’ in the networks (class C). The correlation we observe between gene conservation and local network topology may be due to specific selection pressures, for instance on essential genes^12^.

**Figure 2.**
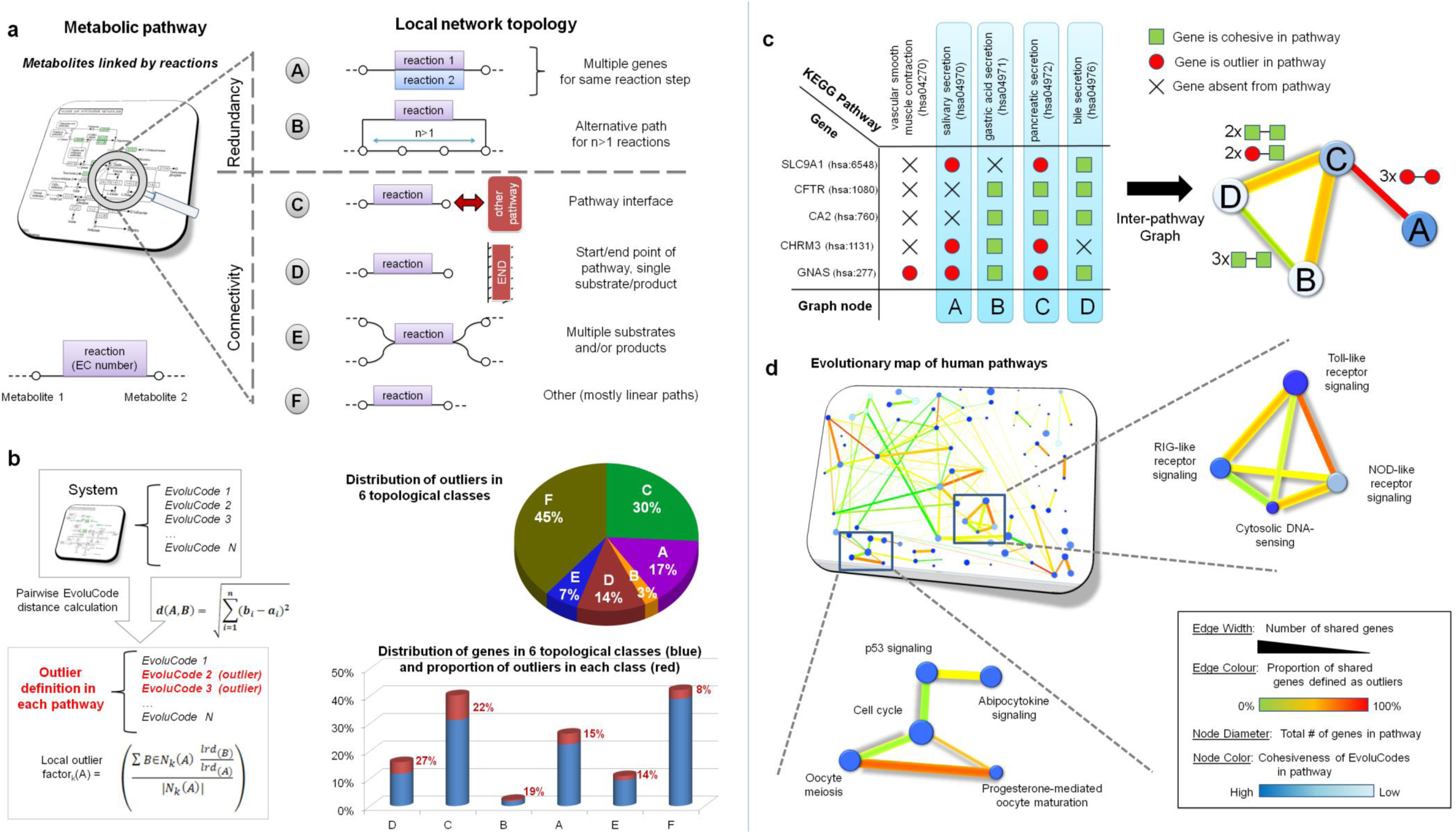
Characterization of outlier genes at the system and cellular levels. (a) Definition of 6 classes of local topological motifs in metabolic pathways, depending on the redundancy and connectivity of the reactions (and associated genes) in the network. (b) Identification of outlier genes and their distribution in the local topology classes. (c) The crosstalk between 2 systems is characterized by the proportion of shared outlier genes, indicated by a color gradient from green (all cohesive) to red (all outlier). (d) An integrated evolutionary map of selected human pathways showing the number and cohesiveness of the gene evolutionary histories, associated with individual pathways (nodes) and pathway crosstalk (edges).

Having established the evolutionary cohesiveness of individual pathways, we then asked whether we could identify unusual evolutionary behavior at the cellular level. Individual pathways often function in a coordinated fashion and understanding the interactions or crosstalk between pathways is important for deciphering complex cellular processes, such as the appropriate physiological responses to internal or external stimuli. To investigate these high-level processes, we identified a set of genes involved in the crosstalk between 155 KEGG pathway maps, reflected by the fact that all the genes in the set were present in at least 3 maps. In this case, the evolutionary cohesiveness of a gene is context-dependent, i.e. a gene may be defined as cohesive in one of these pathways and as an outlier in another (Fig. 2c). Such cases of differential evolutionary conservation may indicate important events, such as gene duplications, rearrangements or losses and the subsequent gain or loss of interactions in the network. For each pair of KEGG maps, we calculated the proportion of outlier genes observed in the overlapping set of genes shared between the two systems. We then constructed a global map of the relationships between the 155 maps, representing the evolutionary behavior of these pathways during vertebrate evolution (Fig. 2d and Supplementary Fig. 4). The exploration of this map provides a powerful and visual means of highlighting important events in the evolution of human biological systems.

Two examples are highlighted in Fig. 2d. First, the genes involved in both cell cycle and oocyte meiosis pathways are generally cohesive with the other genes in these pathways, but the crosstalk with the progesterone-mediated oocyte maturation pathway contains a higher proportion of outlier genes (Supplementary Table 2). In fact, cell cycle and oocyte meiosis pathways are conserved in most vertebrates, while the exact nature of oocyte maturation is more variable between species. A number of these functional specificities are highlighted by the EvoKEN outliers, such as the Myt1 gene coding for a cdc2-inhibitory kinase (PMYT1_HUMAN), which acts as a negative regulator of entry into mitosis during the cell cycle. Inspection of the Myt1 evolutionary barcode (Supplementary Fig. 5a) indicates a more divergent sequence family than is typical for this conserved pathway. This might be a result of the different functions of Myt1, which is implicated in control of entry into meiosis, either alone (as in *Xenopus*) or in concert with Wee1 (as in mouse oocytes)^13^. Other examples of outliers are provided in Supplementary Fig. 5.

The second example concerns the innate immune system, where pattern recognition receptors, such as Toll-like receptors (TLR), RIG-I-like receptors (RLR) or NOD-like receptors (NLR), recognize a wide variety of pathogens and endogenous molecules and trigger complex, overlapping intracellular signaling cascades. Outlier genes involved in the crosstalk between these pathways are described in Supplementary Table 3. We highlight one example: the receptor interacting protein RIP1 (RIPK1_HUMAN), which plays a crucial role in the cellular response to TLR and RLR signals, switching between cell survival through RIP1 activation of NF-κB and cell death induced by caspase-8 cleavage of RIP1^14^. The RIP1 evolutionary barcode (Supplementary Fig. 6a) shows a typical sequence conservation in vertebrate evolution, but synteny is only observed in mammals and not in fishes for example where RIP1 plays a different role in TLR signaling^15^. Other examples of outliers are provided in Supplementary Fig. 6. Unraveling the evolutionary history of these pathways and their crosstalk will be important in understanding how the immune system functions and in developing effective therapeutic and vaccine strategies.

It is clear that more in-depth analysis, involving phylogenetic tree and ancestor reconstruction would be required to describe in detail the evolutionary events identified in these studies. The advantage of EvoKEN is that it provides an effective framework for investigating the evolution of large systems at different granularity levels from local network motifs to the cellular level, allowing the rapid identification of interesting patterns in a particular biological context. Hopefully, EvoKEN will contribute to the emerging field of evolutionary systems biology, with the goal of understanding and modeling the topological and dynamic properties of the complex networks that govern the behavior of the cell.

## METHODS

### Construction of EvoluCodes for the human proteome

The evolutionary barcodes (EvoluCodes) used in this study were constructed as described in ^8^. Each protein-coding human gene is thus associated with one EvoluCode that is visualized as a 2D matrix. The columns of the matrix correspond to the studied organisms, which in this work consist of 17 vertebrates with almost complete genomes from the Ensembl ^16^ database (version 51). The rows of the matrix correspond to different evolutionary parameters Table 1) that were extracted from multiple alignments ^17,18^, synteny analysis and orthology data^19^. For each vertebrate organism, the most closely related homolog to the human reference gene was identified (based on percent residue identity) and 10 parameters were calculated.

**Table 1.**
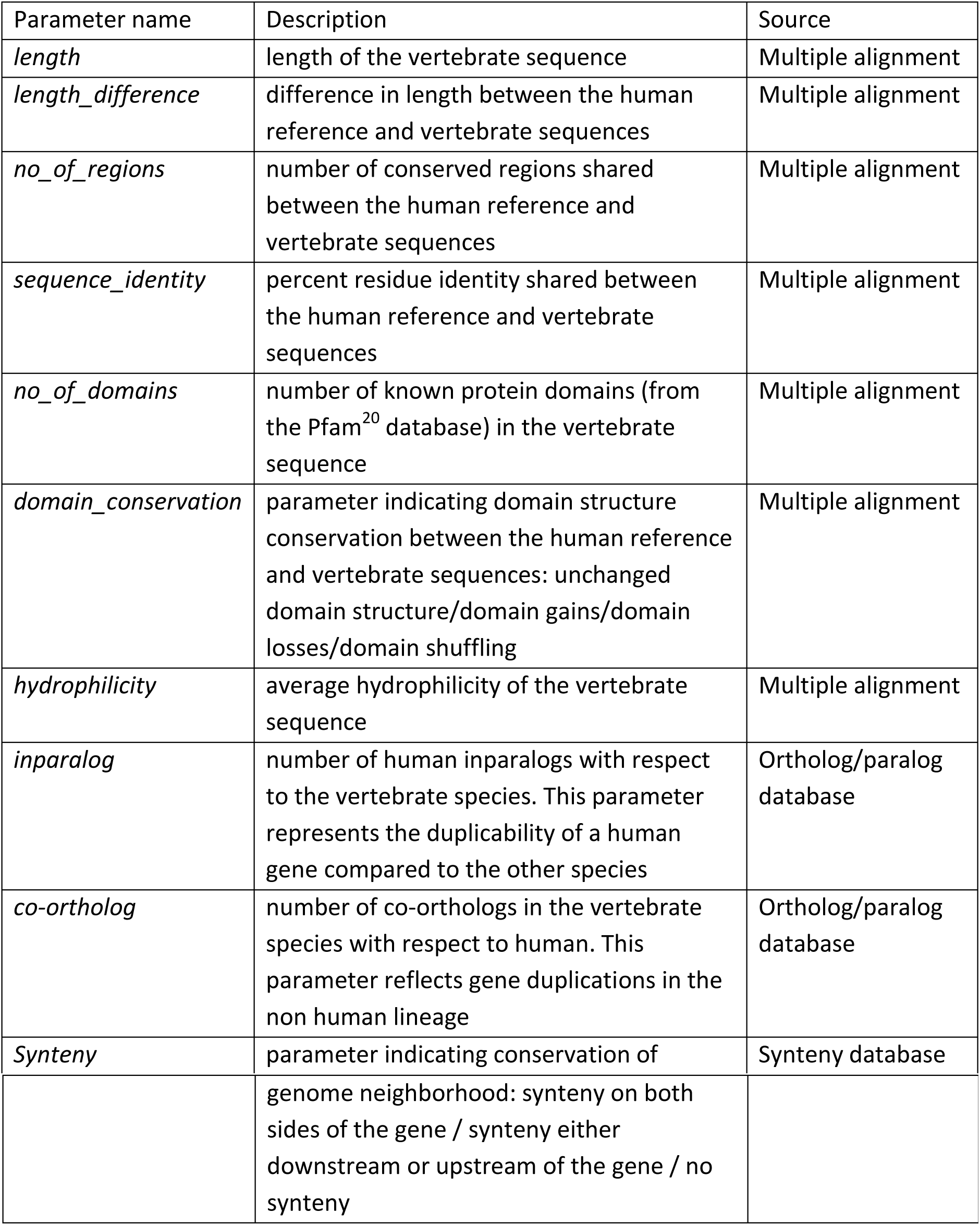
Evolutionary parameters included in EvoluCodes of human proteome.

To facilitate visualization of the EvoluCode, a color is assigned to each matrix cell representing typical or atypical parameter values. To do this, the distribution of each parameter in each organism is first described by the sample percentiles, using the Emerson-Strenio formulas ^21^ implemented in the R software and color gradients are assigned to three intervals:

- Interval 1 represents values that are lower than what is generally observed for a specific parameter in a specific organism and is assigned a blue-to-green gradient
- Interval 2 represents values that correspond to what is generally observed for a specific parameter in a specific organism and is assigned a green color
- Interval 3 represents values that are higher than what is generally observed for a specific parameter in a specific organism and is assigned a green-to-red gradient.

By compiling several evolutionary parameters extracted from different biological levels, from residue data to phylum data, EvoluCodes incorporate an evolutionary systems biology point of view. Consequently, EvoluCodes can highlight important evolutionary events that could not be discovered using a single evolutionary parameter such as sequence conservation or domain composition. The complete set of 19778 human EvoluCodes can be visualized online at: lbgi.igbmc.fr/barcodes.

### Analysis of human pathway data and definition of cohesive/outlier genes

We based our analysis on pathway data from the Kyoto Encyclopedia of Genes and Genomes (KEGG) knowledge base. We analyzed 248 human pathways with the help of the KEGG SOAP server (http://www.kegg.jp/kegg/soap/). A total of 5849 EvoluCodes could be mapped to the genes in these pathways.

For each pathway, we then identified ‘outlier’ genes, i.e. genes with an unusual evolutionary history (EvoluCode) compared to the other genes in the pathway. We determined outliers using an anomaly detection algorithm called Local Outlier Factor (LOF) ^22^. The basic concept of LOF is the local density, where locality is given by k nearest neighbors. By comparing the local density of an evolutionary barcode to the local densities of its neighbors, we identify regions of similar density, as well as barcodes that have a substantially lower density than their neighbors. These are considered to be outliers. The local density is estimated by the typical distance at which a barcode can be “reached” from its neighbors.

First, the 2D matrix representing an EvoluCode, consisting of 10 rows and 17 columns, is redimensioned to a 1D vector of length, n=170. Then, if A and B are 2 EvoluCodes in Euclidean n-space, with A = (a_1_, a_2_,…,a_n_) and B = (b_1_, b_2_,…,b_n_), the distance between A and B is:

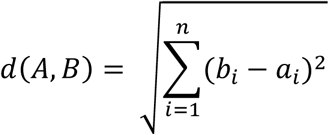

The Euclidean distance between the EvoluCode A and its k nearest neighbors is denoted kdist(A) and the set of k nearest neighbors is N_k_(A). The reachability distance is then calculated as:

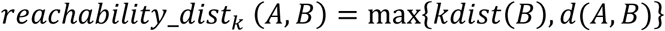

The local reachability density (lrd) of EvoluCode A is defined as:

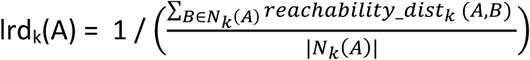

And the local reachability densities are then compared with those of the neighbors using:

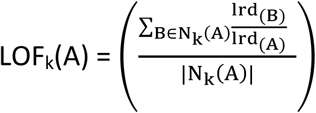

The LOF score thus represents the cohesiveness of the EvoluCode associated with each gene in the context of its pathway. The authors of the LOF algorithm consider that a score less than 1 indicates a clear inlier object, i.e. a cohesive barcode. Genes with a LOF score significantly greater than 1 are considered as outliers. However, the threshold determining a clear outlier depends on the dataset. Here, we defined the outlier threshold value as the upper quartile for the LOF scores of the EvoluCodes in the context of the 248 human pathways, which was 1.037.

### Analysis of metabolic pathways

The KEGG database currently contains pathway data for 84 human KEGG metabolic pathways, where the nodes in the networks represent metabolites (substrates, intermediates and products). The edges between nodes represent reactions that are associated with one or more genes/proteins. For our experiment, we selected all pathways with more than 20 human genes, giving us an initial set of 20 pathways, containing a total of 875 different reactions, of which 671 reactions were associated with cohesive genes and 204 reactions with outlier genes. We defined 6 classes of local topological motifs within these pathways, based on node redundancy and connectivity (Table 2).

**Table 2.** Definition of local network topology classes.

| Class | Redundancy | Connectivity | Description |
| --- | --- | --- | --- |
| A | yes | N/A | Alternative gene for same reaction step |
| B | yes | N/A | Alternative path for $n > 1$ reactions |
| C | N/A | Inter-pathway | Pathway interface |
| D | N/A | 1 intra-pathway | Start/end of pathway, single substrate/product |
| E | N/A | $> 2$ intra-pathway | Multiple substrates and/or products |
| F | No | 2 intra-pathway | Other: mostly linear paths, plus a small number of exceptions, such as unlinked genes |

We then determined the topological localization of all biological reactions associated with the outlier genes. For each class, we calculated the proportion of cases where the reaction was associated with an outlier gene. In cases where a reaction was associated with more than one gene (protein complexes, genes with similar biochemical functions, etc.), we used the gene with the lowest LOF score. This choice reduced the number of reactions that are considered as outliers and only reactions with a clear outlier status were included for analysis.

### Construction of the cellular level evolutionary map

We constructed a cellular level map, representing the evolutionary histories of the pathways in the KEGG database. For the 200 human KEGG pathways, we identified the genes shared by each pair of pathways. We then focused our analysis on the pathway pairs sharing at least 3 genes, representing 155 KEGG pathways, which describe mainly cellular processes and signal transductions. In the evolutionary map, each node represents a specific KEGG pathway and the edge joining 2 nodes represents the genes shared by the two pathways. The node diameter is proportional to the number of genes implicated in the pathway. Each node is assigned a color representing the homogeneity of the EvoluCodes associated with the genes. The cohesiveness of the pathway evolution is estimated based on LOF value dispersion, using the IQR (interquartile range, IQR = Q_3_ − Q_1_). A low IQR indicates more cohesive barcodes associated with a given node.

Pathways with high cohesiveness are indicated by dark blue and pathways with low cohesiveness are light blue. The edge thickness is proportional to the number of genes shared by the 2 nodes, while the edge color indicates the proportion of shared genes identified as outliers in one or both linked pathways. A green edge links pathways that do not share any outlier genes. A red edge links pathways where all shared genes are outliers in at least one of the maps. Intermediate values are assigned a green to red color gradient.

## SUPPLEMENTARY INFORMATION

Supplementary Figures 1-6 and Supplementary Tables 1-3 are are available as supplementary files.

## ACKNOWLEDGMENTS

This work was supported by the Decrypthon program, co-funded by Association Française contre les Myopathies (AFM, 14390-15392), IBM and Centre National de la Recherche Scientifique (CNRS). We acknowledge financial support from the ANR (EvolHHuPro: BLAN07-1-198915 and Puzzle-Fit: 09-PIRI-0018-02) and Institute funds from the CNRS, INSERM, and the Université de Strasbourg.

## AUTHOR CONTRIBUTIONS

B.L., J.D.T. and O.P. conceived the study. B.L. and N.H.N. devised and implemented the algorithm and conducted experiments. B.L., N.H.N. O.L. J.D.T. and O.P. discussed the results and implications. O.L. and O.P. evaluated the biological relevance of the results. J.D.T. supervised the project. B.L. and J.D.T. wrote the manuscript.

## REFERENCES

1. T. F. Mackay, E. A. Stone, and J. F. Ayroles, Nat Rev Genet 10 (8), 565 (2009).

2. A. L. Barabasi, N. Gulbahce, and J. Loscalzo, Nat Rev Genet 12 (1), 56 (2011).

3. A. Schuler and E. Bornberg-Bauer, Methods Mol Biol 696, 273 (2011).

4. T. Yamada and P. Bork, Nat Rev Mol Cell Biol 10 (11), 791 (2009).

5. R. A. Pache and P. Aloy, PLoS One 7 (2), e31220 (2012).

6. G. Musso, A. Emili, and Z. Zhang, Methods Mol Biol 856, 363 (2012).

7. D. F. Veiga, B. Dutta, and G. Balazsi, Mol Biosyst 6 (3), 469 (2010).

8. B. Linard, N. H. Nguyen, F. Prosdocimi et al., Evol Bioinform Online 8, 61 (2012).

9. M. Kanehisa, S. Goto, Y. Sato et al., Nucleic Acids Res 40 (Database issue), D109 (2012).

10. J.H.M. Janssens, I. Flesch, and Postma. E.O., presented at the 8th International Conference on Machine Learning and Applications, Miami, USA, 2009.

11. L. Fokkens and B. Snel, PLoS Comput Biol 5 (1), e1000276 (2009).

12. E. V. Koonin and Y. I. Wolf, Nat Rev Genet 11 (7), 487 (2010).

13. M. Gaffre, A. Martoriati, N. Belhachemi et al., Development 138 (17), 3735 (2011).

14. N. Festjens, T. Vanden Berghe, S. Cornelis et al., Cell Death Differ 14 (3), 400 (2007).

15. A. Rebl, T. Goldammer, and H. M. Seyfert, Vet Immunol Immunopathol 134 (3-4), 139 (2010).

16. P. Flicek, M. R. Amode, D. Barrell et al., Nucleic Acids Res 40 (Database issue), D84 (2012).

17. F. Plewniak, L. Bianchetti, Y. Brelivet et al., Nucleic Acids Res 31 (13), 3829 (2003).

18. J. D. Thompson, A. Muller, A. Waterhouse et al., BMC Bioinformatics 7, 318 (2006).

19. B. Linard, J. D. Thompson, O. Poch et al., BMC Bioinformatics 12, 11 (2011).

20. M. Punta, P. C. Coggill, R. Y. Eberhardt et al., Nucleic Acids Res 40 (Database issue), D290 (2012).

21. J.D. Emerson and Strenio J., in Understanding Robust and Exploratory Data Analysis, edited by J.W. Tukey F. Mosteller, D.C. Hoaglin (Wiley, New York, 1983), pp. 58.

22. H.P. Kriegel, M.M. Breunig, R.T. Ng et al., presented at the ACM SIGMOD Int. Conf., 2000.

